# Chronic cervical vibration ameliorates depression-like behavior in Wistar-Kyoto rats

**DOI:** 10.1101/2025.11.17.688804

**Authors:** Sei-etsu Fujiwara, Sakae Narumi, Chiaki Kakehashi, Hitomi Fujioka

## Abstract

Major depressive disorder (MDD) is a highly prevalent psychiatric condition, and a substantial proportion of patients exhibit inadequate responses to conventional pharmacotherapy. Somatosensory processing abnormalities and altered thalamo-cortical connectivity have been implicated in the pathophysiology of MDD, raising the possibility that targeted somatosensory stimulation may modulate affective circuits. Here, we examined whether chronic cervical vibration improves depression-like behaviors in Wistar-Kyoto (WKY) rats, a well-established model of treatment-resistant depression.

Rats received repeated vibration stimulation applied to the cervical region. Behavioral effects were evaluated using the novelty-suppressed feeding test (NSFT) and the forced swim test (FST). Swimming activity was quantified during the late-phase of the FST to exclude early panic-like behavior. Immunohistochemistry for c-Fos was performed in the dorsal raphe (DR), lateral habenula (LHb), paraventricular nucleus of the hypothalamus (PVN), and prefrontal cortex (PFC).

Vibration-treated rats showed significantly shorter feeding latencies in the NSFT and significantly increased swimming distance during the analysis period of the FST. No significant group differences in c-Fos expression were observed in DR, LHb, PVN, or PFC.

These findings indicate that chronic cervical vibration produces reproducible improvements in depressive-like behavior in WKY rats. Because c-Fos activity remained unchanged, the behavioral effects may reflect gradual changes in broader network processing rather than acute activation of monoaminergic or habenular-prefrontal pathways. Cervical vibration represents a simple, non-invasive sensory intervention with translational potential. Additional mechanistic studies are warranted.

## 1. Introduction

Major depressive disorder (MDD) is one of the most prevalent psychiatric illnesses, and despite available treatments, approximately one-third of patients fail to achieve remission [1]. This highlights the need for new therapeutic approaches that are safe, affordable, and effective for treatment-resistant depression.

Growing evidence suggests that abnormalities in somatosensory pathways contribute to depressive symptomatology. MDD patients show enhanced thalamus–primary somatosensory cortex (SI) connectivity [2], prolonged somatosensory evoked potential latency [3], increased medial thalamus–somatosensory/temporal connectivity [4], and region-dependent disruptions in thalamo-cortical coupling [5]. These findings implicate altered sensory integration in the neurobiological basis of depression.

Somatosensory stimulation itself has shown promise as a neuromodulatory intervention. A clinical study reported beneficial effects of peripheral somatosensory stimulation on depressive and anxiety symptoms. In rodent models, whole-body vibration attenuated depression-like behaviors induced by chronic stress [10] and improved anxiety-like behavior, cognition, and hippocampal neuroinflammation in aged rats [11]. Local peripheral vibration can also influence monoaminergic signaling in the brain [12], suggesting direct access to affect-related neuromodulatory pathways.

The cerebellum has emerged as a key regulator of affective processes. Enhancing cerebellar cortical activity improves depression-like behavior, while inhibition produces depressive phenotypes [6]. Anatomical studies show robust cerebello-thalamo-prefrontal projections [7], and the PFC sends direct projections to the lateral habenula (LHb) [8], a major node in aversive processing. Importantly, somatosensory input from trunk and neck regions projects to cerebellar areas closely aligned with affective domains [9].

Wistar-Kyoto (WKY) rats exhibit high anxiety, increased behavioral despair, and poor antidepressant responsiveness [13], making them a suitable model of treatment-resistant depression. Whether chronic peripheral neck vibration can improve depression-like behaviors in this strain remains unknown.

To address this question, we investigated the effects of chronic cervical vibration on behavior and neural activity in WKY rats using NSFT, late-phase FST, and c-Fos immunohistochemistry.

## 2. Materials and Methods

### 2.1 Animals

Male WKY rats (9–14 weeks old; Jackson Laboratory Japan) were housed individually under controlled temperature (23 ± 2°C), humidity (60 ± 5%), and a 12:12 h light–dark cycle. Food and water were available ad libitum. After a 1-week acclimation period, rats were assigned to either the vibration or control group. All experimental procedures were approved by the Animal Experiments Committee of St. Marianna University School of Medicine and were conducted in accordance with institutional guidelines.

### 2.2 Vibration device

A vibration motor with an eccentric rotating mass was embedded within a plastic cylindrical housing. A single-cell AA battery case was affixed using epoxy resin, providing continuous operation for more than 30 min.

Because this device is the subject of an ongoing patent application, certain proprietary engineering details—including internal circuitry, precise frequency–acceleration characteristics, and component-level specifications—are not disclosed in this Methods section to protect intellectual property rights. All non-proprietary information necessary for evaluating the scientific validity of the study is fully provided.

### 2.3 Cervical vibration protocol

Rats were fitted with a rat jacket (Bio Research Center), to which the vibration device was secured over the cervical region. Vibration stimulation was delivered for 30 min per session, across multiple sessions over several weeks, during the light phase. Control animals wore an identical device with the vibration function disabled. Throughout the study period, no abnormal behaviors, dermatitis, wounds, weight loss, or attempts to remove the device were observed. Animals maintained normal grooming and exploratory behaviors. In alignment with intellectual property protection measures associated with the device (patent application pending), specific operational parameters—such as the exact stimulation waveform, internal control architecture, and proprietary tuning procedures—cannot be disclosed. However, the overall stimulation schedule and experimental workflow are reported in full to maintain methodological transparency.

### 2.4 Novelty-Suppressed Feeding Test (NSFT)

Rats were food-deprived for 22–24 h prior to testing. The test was conducted in a novel cage containing clean bedding and a single food pellet. Latency to bite the pellet was recorded via video and directional microphone. After testing, rats were returned to their home cage and allowed to feed ad libitum.

### 2.5 Forced Swim Test (FST)

Rats were placed in a cylindrical tank filled to a depth of 30 cm with water maintained at 23– 25°C for 300 s. Behavior was video-recorded. As escape-related activity during the initial 150 s is considered non-specific, analyses focused on the 150-300 s interval. Locomotor activity was quantified as total distance traveled.

### 2.6 Immunohistochemistry

Two hours after the final vibration or control session, rats were perfused transcardially with 4% formaldehyde. Brains were post-fixed, cryoprotected, and sectioned at 30 *μ*m. Free-floating sections containing the DR, LHb, PVN, or PFC were incubated with primary antibodies (c-Fos, TPH2) followed by appropriate fluorescent secondary antibodies. Sections were imaged using fluorescence and confocal microscopy. Regions of interest were delineated using NeuroTrace or TPH2 labeling, and c-Fos-positive cells were quantified using SimpleCellCounter.

### 2.7 Statistical analysis

All statistical analyses were performed using GraphPad Prism 10.4.0 for immunohistochemical data and custom-written LabVIEW code for behavioral data. Data are presented as mean ± SEM. Behavioral data were analyzed using Welch’s t-test, with degrees of freedom calculated according to the Welch–Satterthwaite approximation. Statistical significance was defined as p < 0.05 (two-tailed). No data points were excluded from the analyses.

## 3. Results

### 3.1 Chronic cervical vibration significantly reduced feeding latency in the NSFT

Vibration-treated rats showed markedly shorter feeding latency (Figure 1A)

**Figure 1:**
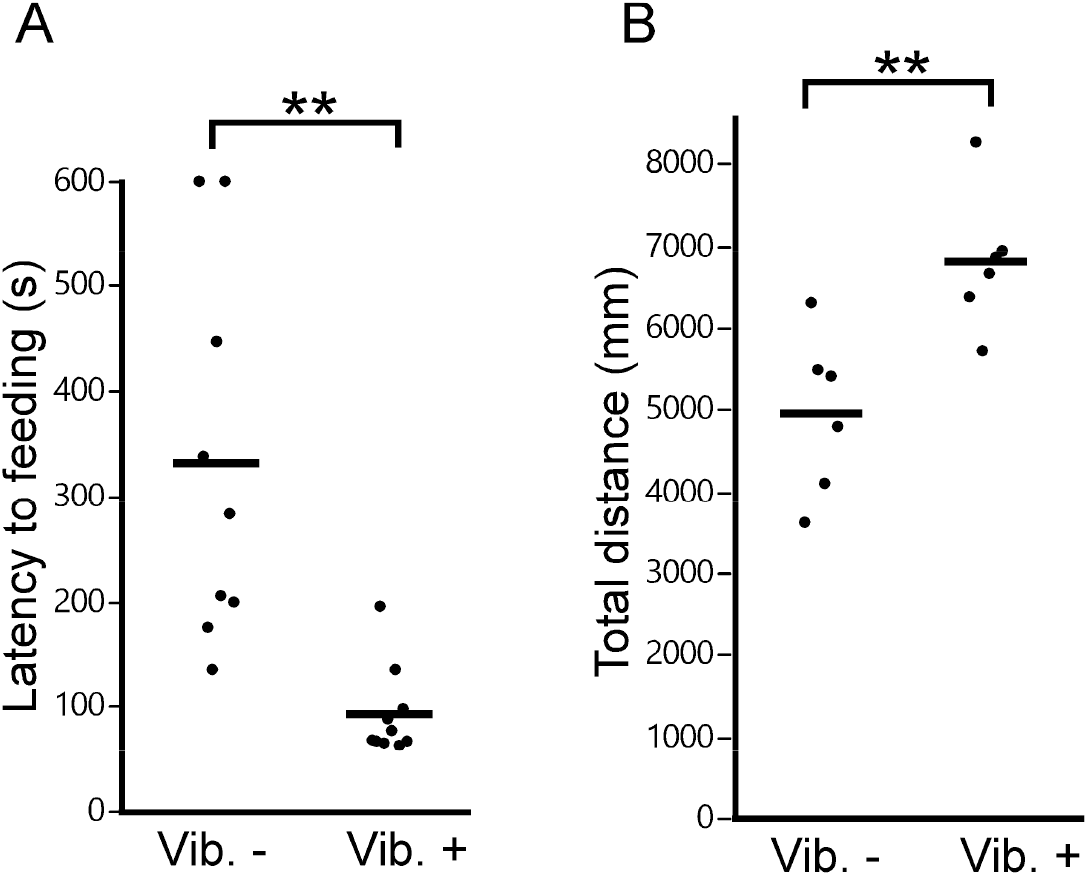
Chronic cervical vibration improves depression-like behaviors in WKY rats. (A) Feeding latency in the NSFT. Vibration significantly reduced latency (Welch’s t-test: t(10.0)=3.7808, p=0.0030). (B) Total distance traveled during the 5–10 min period of the FST. Vibration significantly increased swimming activity (t(7.93)=–3.5000, p=0.00594). Dots: individual rats; bars: group means.

### 3.2 Chronic cervical vibration increased locomotor activity in the FST

Total distance traveled during the 5–10 min period was significantly greater in vibration-treated rats (Figure 1B).

### 3.3 c-Fos expression did not differ between groups

No significant differences were observed in c-Fos–positive cell counts in DR, LHb, PVN, or PFC (all p > 0.05)(Data not shown).

## 4. Discussion

This study demonstrates that chronic cervical vibration significantly improves depressive-like behavior in WKY rats. Vibration reduced feeding latency in the NSFT and enhanced swimming activity during the late-phase (5–10 min) FST. Because the early panic-like period was excluded, the FST results reflect sustained behavioral state rather than acute arousal.

The beneficial behavioral effects observed here are consistent with previous reports that somatosensory stimulation can modulate mood-related pathways. Peripheral somatosensory stimulation has shown clinical benefit for depression and anxiety. Whole-body vibration reduces depression-like and anxiety-like behaviors in rodent models [10,11], while local vibration influences monoaminergic systems [12]. These studies support the plausibility that cervical vibration can alter neural circuits relevant to affect.

Although we found no significant c-Fos differences in the DR, LHb, PVN, or PFC, this does not preclude slower or circuit-level mechanisms. Somatosensory inputs projecting to the cerebellum [9] may modulate cerebello-thalamo-prefrontal circuits implicated in emotional regulation [6– 8]. Altered thalamo-cortical connectivity—well-documented in MDD patients [2–5]—may also be responsive to repeated neck-derived mechanosensory input.

The WKY strain exhibits characteristic depressive- and anxiety-like behaviors and limited antidepressant responsiveness [13], making the behavioral improvements seen here particularly meaningful for treatment-resistant depression research.

Future studies incorporating electrophysiology, functional connectivity mapping, and molecular markers will be needed to determine how cervical vibration produces lasting behavioral improvements.

Chronic cervical vibration is non-invasive, low-cost, and technically simple, suggesting potential as a novel sensory neuromodulatory strategy.

## Acknowledgments

This study was supported by JSPS Kakenhi 22H04922, 22H03631, 23K24887, 24K15868, 24K11729, 24H01435, 25K15206, Tateishi Science and Technology Foundation, and “Innovation inspired by Nature” Research Support Program, SEKISUI CHEMICAL CO., LTD.

## Animal ethics

All procedures performed in studies involving animal experiments were approved by the Animal Experiments Committee of St. Marianna University School of Medicine and were in accordance with their guidelines.

